# Within-colony color variation in a chalice coral is associated with region-specific pigment expression and higher photoconvertible pigment transcripts

**DOI:** 10.64898/2026.08.29.746859

**Authors:** Oskar Baumgarte, Zane J. Moore

## Abstract

A single colony of coral often demonstrates conspicuous color variation. However, the molecular basis of this variation within one clonal individual is poorly understood. While coral tissues within a colony have distinct colors, they share the same genetic background, suggesting differences between them are regulatory rather than genetic. This pilot study aims to characterize the transcriptional mechanisms of color regions in a green-and-orange chalice coral colony through an end-to-end transcriptomic analysis. First, we sampled the green body, an orange oral region, and a transition region between them. Then, a holobiont transcriptome was assembled de novo from paired-end RNA-seq of each color region. In the holobiont transcriptome, coral host transcripts were separated from dinoflagellate symbiont transcripts. Green fluorescent protein (GFP)-like pigment genes were classified phylogenetically. Their expression was then quantified using host-only normalization, which corrects for a large regional difference in symbiont read fraction, and compared across color regions. Relative to the green body, the orange oral region showed higher expression of two sets of host pigment genes: distinct orange fluorescent-protein and chromoprotein transcripts, and green-to-red photoconvertible EosFP-family proteins whose unconverted state is green, implicating photoconversion as a contributing mechanism. This study reveals that distinct color regions within a single colony are associated with region-specific up-regulation of different pigment transcripts, including both directly colored proteins and photoconvertible proteins. Further research is needed to confirm spectrally that photoconversion contributes to the coloration and test whether the pattern holds across multiple colonies.

## Introduction

Reef corals are among the most vividly pigmented animals in the sea. Much of their color comes from a family of green fluorescent proteins (GFPs). This family includes cyan, green, and red fluorescent proteins and non-fluorescent chromoproteins [1, 2]. These pigments are encoded by the nuclear genome of the coral, not by its algal symbionts [3]. Because the host nuclear genome carries these genes, the coral itself controls where and how strongly they are expressed, making this pigmentation a property of host pigment expression rather than of algal pigment content [3, 4]. These pigment genes have been linked to photoprotection and photosynthetic light management [4–6], antioxidant activity [7], and prey attraction [8], with their expression varying with environmental conditions and developmental stage. Within a single colony, these same genes can produce different colors in adjacent tissue, and how this occurs is not well understood.

In corals, the observed pigmentation belongs to a holobiont, which refers to the coral host together with the intracellular dinoflagellate symbionts of the family Symbiodiniaceae. This makes interpreting coral color even more complex because both the coral host and its symbionts contribute to observed color. Moreover, symbiont density strongly affects apparent color, and it has been found that browner, symbiont-dense tissue can mask differences in host pigment [3]. Therefore, separating host pigment-gene regulation from variation in symbiont density becomes essential.

Some GFP-like proteins undergo photoconversion, an irreversible change in emission color triggered by light [9, 10]. Specifically, photoconvertible members of the EosFP and Kaede families are synthesized in a green-emitting state. EosFP was isolated from the lobophylliid coral *Lobophyllia hemprichii* and Kaede from *Trachyphyllia geoffroyi* [9, 10]. Both belong to the GFP-like superfamily and share the His-Tyr-Gly chromophore that undergoes green-to-red conversion, and coral FP genes of this family occur as multicopy arrays rather than as single loci [11]. Upon the absorption of near-UV or violet light, they undergo an irreversible green-to-red conversion [9, 10]. Photoconversion has also been proposed as a route to orange-red coloration in corals [12].

As the initial state of a photoconvertible protein is green, genetic classification categorizes it as green even though it can demonstrate the red color. Thus, the color class inferred from sequence phylogeny may not match the tissue color. This discrepancy can be tested directly. If phylogenetic color class describes the color a transcript contributes, then class should predict the direction of change between regions: green-classified transcripts should be more abundant in the green body and less abundant in the orange region, with the reverse for orange- and red-classified transcripts. In this study, we use direction to indicate whether a transcript is more abundant in the orange region or the green body.

Intra-colony color variation, where one colony carries two or more distinct colors, offers a way to test this directly. Here, because all regions of the colony share the same genetic background, color differences between regions are unlikely to arise from differences in gene copy number or allelic identity, and are instead attributable to differences in regulation, splicing, or post-translational processing of a shared host nuclear genome. A single colony is the closest approach to a common genetic background that is feasible in a coral, with the only expected genotypic differences being rare somatic mutations accumulated during colony growth.

Here we examine one chalice coral colony of *Echinophyllia* sp., shown in Figure 1. We use region to refer to a visually distinct area of the colony surface defined by color. The colony has three visually distinct regions, referred to as CG (the green body), CI (the intermediate transition region), and CO (the orange oral region).

**Figure 1.**
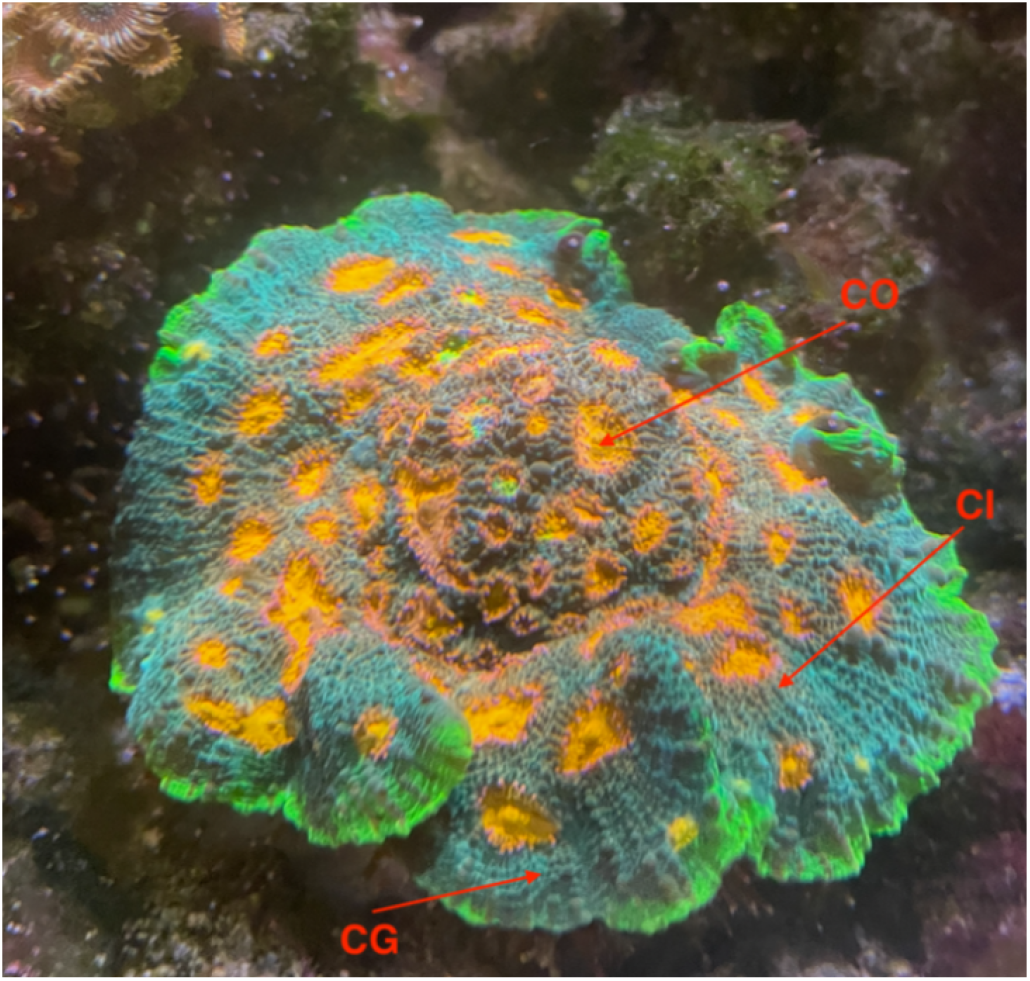
This single colony of *Echinophyllia* sp. displays three color regions: a green body (CG), an orange oral region (CO), and a transition region (CI) where the color grades from green to orange between them.

We evaluated four non-exclusive hypotheses for the mechanism producing distinct color regions:

1. Pigment transcripts differ in abundance between color regions as predicted by their color class: GFP and CFP transcripts are more abundant in CG, and OFP and RFP transcripts are more abundant in CO.
2. Distinct pigment transcripts, rather than uniform change across the same set of transcripts, distinguish the regions.
3. The symbiont share of the transcriptome covaries with color region rather than with host pigment expression.
4. The color of the tissue is not fully determined by sequence: post-translational change of state, such as green-to-red photoconversion, contributes to the orange coloration.

## Methods

Figure 2 summarizes the full workflow of this study.

**Figure 2.**
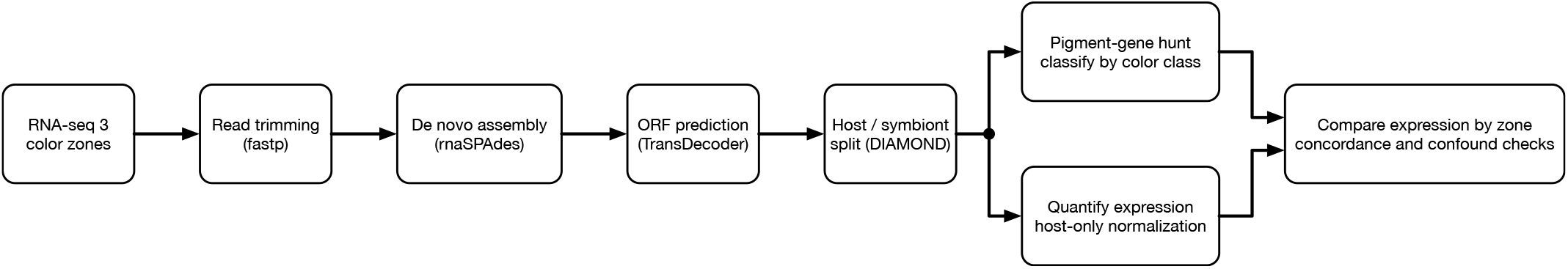
End-to-end coral holobiont transcriptomics workflow. RNA-seq data from three color regions (CO, CI, CG) were trimmed and quality-filtered, assembled de novo via rnaSPAdes, and resolved into host and symbiont transcripts. Host-normalized expression was subsequently quantified across color regions, and expression patterns were compared by region concordance while accounting for potential confounding effects from regional differences in symbiont abundance.

### Sampling and RNA sequencing

Three samples of coral tissue were collected from the colony, each corresponding to the CG region, the CI region, and the CO region, by scraping with an X-Acto knife. Total RNA was extracted from each sample. Paired-end RNA sequencing (RNA-seq) libraries were prepared and sequenced by Novo-gene, with one library for each region. The integrity of the raw FASTQ data was verified against provider MD5 check-sums.

### Quality control and read trimming

Adapters and low-quality tails were removed from the reads using paired-end auto-detection with quality and length filtering in fastp [13]. Using FastQC and MultiQC, read quality was assessed before and after trimming. Per-sample read retention ranged from 99.4% to 99.5%, with 99.45% retained overall.

### De novo holobiont assembly

Trimmed reads from all three regions were pooled and assembled de novo using rnaSPAdes [14]. The assembly contained 329,871 transcripts and approximately 322 Mbp, with an N50 of about 1,900 bp and 46.4% GC content. As all tissues came from one genotype, a single pooled assembly was appropriate. De novo assembly was initially attempted with Trinity but could not be completed within the available computational resources [15]. Therefore, rnaSPAdes was used instead, as it is a standard, well-benchmarked alternative.

### Host and symbiont deconvolution

Using TransDecoder, a single open reading frame per transcript was predicted [16]. DIAMOND blastp was used to classify predicted proteins against cnidarian host protein sets and Symbiodiniaceae protein sets, keeping the best match at an e-value threshold of 1e-5 [17]. The cnidarian sets included *Acropora, Orbicella, Nematostella*, and *Exaiptasia*. The Symbiodiniaceae sets included *Symbiodinium, Breviolum, Cladocopium*, and *Durusdinium*. Instead of a symmetric best-bitscore rule, host-conservative assignment was applied: a transcript was labeled symbiont only when its best symbiont bitscore reached ≥ 45 bits and exceeded its best host bitscore by more than 5 bits. A transcript with any qualifying match to the host set was labeled host, and transcripts with no host hit and no symbiont hit meeting the bitscore criteria were marked as unresolved. Among the transcripts with predicted ORFs, 49,854 were host-assigned, 71,090 were symbiont-assigned, and 14,537 were marked as unresolved. For downstream analysis, a host-only transcriptome of 64,391 transcripts, including the 49,854 host-assigned transcripts added to the 14,537 unresolved transcripts, was kept. The unresolved transcripts were retained because almost all of them (14,536 of 14,537) returned no symbiont match and are therefore unlikely to be dinoflagellate in origin, while many of the coral genes are missing close database matches and would be lost by removing no-hit transcripts. Pigment candidates were verified against the NCBI non-redundant database to confirm cnidarian rather than dinoflagellate origin.

### Pigment-gene identification and phylogenetic color classification

A reference set of 892 GFP-like proteins was obtained from FPbase [18]. It includes anthozoan and hydrozoan fluorescent proteins, chromoproteins, and engineered variants. Candidate color classes were assigned using the framework of Alieva et al. [2]. The Alieva scheme distinguishes cyan, green, and red fluorescent proteins and chromoproteins; OFP and YFP were assigned additionally from the emission maximum annotated for the nearest reference protein in FPbase [18]. Using tblastn, the candidate pigment transcripts were found by searching these references against the host transcriptome. Potential candidates and a 226-protein backbone were aligned with MAFFT [19] and put onto a maximum-likelihood tree with IQ-TREE [20]. From the tree, each candidate was assigned a phylogenetic color class. Specifically, each candidate was assigned as cyan fluorescent protein (CFP), green fluorescent protein (GFP), yellow fluorescent protein (YFP), orange fluorescent protein (OFP), red fluorescent protein (RFP), or chromoprotein. Photoconvertible sequences of the EosFP or Kaede family were also assigned. Photoconvertible proteins are green before they convert, so phylogeny classifies them as green even though they can emit red. Photoconvertibility and emission color were taken from the FPbase annotation for each reference protein [18]. Candidates whose nearest reference was annotated as photoconvertible were flagged so they could be reexamined in later analyses.

### Quantification and host-only-normalized expression

In selective-alignment mode in Salmon, transcript abundances were estimated and mapped against the full holobiont assembly for all three groups [21, 22]. There were 20.6 million CG reads, 22.4 million CI reads, and 23.4 million CO reads. Abundances were imported with tximport [23]. The regions went from 68% symbiont reads in the green body to 32% in the orange region, and this difference would distort normalization if left uncorrected. To prevent this, DESeq2 size factors were computed using only the host transcripts, which gave host-normalized counts [24]. Normalized counts, transcripts per million (TPM), and descriptive log2 fold-changes were reported for CO versus CG, CI versus CG, and CO versus CI.

### Direction-concordance test against the color-class prediction

The test evaluates whether the phylogenetic color class of a pigment gene predicts the region in which it peaks. GFP and CFP are predicted to decline from the green body to the orange oral region, and OFP and RFP are predicted to rise. Chromoproteins were not given a directional prediction: chromoprotein is a functional, non-fluorescent class rather than a color, so no green-to-orange direction follows from it, and they were excluded from the test. YFP was also excluded because yellow is intermediate between green and orange and yields no directional prediction. For each candidate, the direction was calculated as the sign of CO minus CG, and it was labeled concordant if it matched the class prediction. Candidates without a directional class prediction, or with exactly equal endpoints, were removed. Significance was assessed with a label-permutation test. 50,000 permutations were run with a fixed random seed. In each, color-class predictions were shuffled randomly among the transcripts while the observed directions were fixed, and the number of concordant transcripts was counted. This gives the concordance expected by chance, against which the real result was compared. The p-value was calculated as the proportion of permutations in which the number of concordant transcripts equalled or exceeded the observed count. This test is computed across transcripts and is useful despite n = 1 per region. As a secondary check, the same label-permutation test was applied to the subset of strictly monotonic candidates. A candidate is strictly monotonic if its expression changes consistently in one direction across all three regions, either steadily increasing (CG *<* CI *<* CO) or steadily decreasing (CG *>* CI *>* CO), with no reversal in between. Restricting to this subset gives a null that accounts for the same class-label imbalance. The original concordance test was then repeated on three subsets: the top 10 and top 50 transcripts by peak expression, and transcripts above 1,000 normalized counts. This checks whether the abundant transcripts behave as their class predicts, even if the whole list does not.

### Identity analysis of green-classified transcripts increasing toward the oral region

Some green-classified candidates rose from CG to CO, contrary to the predicted decline for that class. For each of these, we recorded the nearest reference protein, along with the emission color and photoconvertibility annotated for that reference in FPbase [18].

### Symbiont-fraction analysis

The per-region symbiont read fraction was computed as the proportion of mapped reads assigned to symbiont transcripts relative to all mapped reads, including those assigned to unresolved transcripts. Partitioning holobiont RNA-seq into host and symbiont fractions is established practice [25], although the resulting fraction reflects the symbiont contribution to the holobiont transcriptome rather than symbiont cell density directly, since it depends on both the number of symbiont cells and their per-cell transcriptional output. This measure was used to test whether regional differences in symbiont read fraction could account for the color difference, as proposed in Hypothesis 3, and to check whether the host pigment signal persisted after host-only size factors were applied.

## Results

Orange fluorescent-protein and chromoprotein transcripts were more abundant in the orange oral region than in the green body. The higher abundance of orange fluorescent-protein transcripts followed the direction predicted by their color class. We expected green-classified transcripts to decline toward the orange region, but they did not. Most were more abundant in CO than in CG. Phylogenetic color class therefore did not predict which region a pigment transcript peaked in. Of 166 pigment candidates, directional class prediction was able to be tested on 102 candidates. Only 27 of the 102 followed the expected direction of change, which is 26.5%. This ratio is at the chance expectation of about 26% (Table 3). A label-permutation test, in which color-class labels were shuffled across transcripts while observed directions were held fixed, gave p = 0.59, demonstrating no convincing evidence that phylogenetic color class predicts the direction of expression change across color regions (Figure 3).

**Figure 3.**
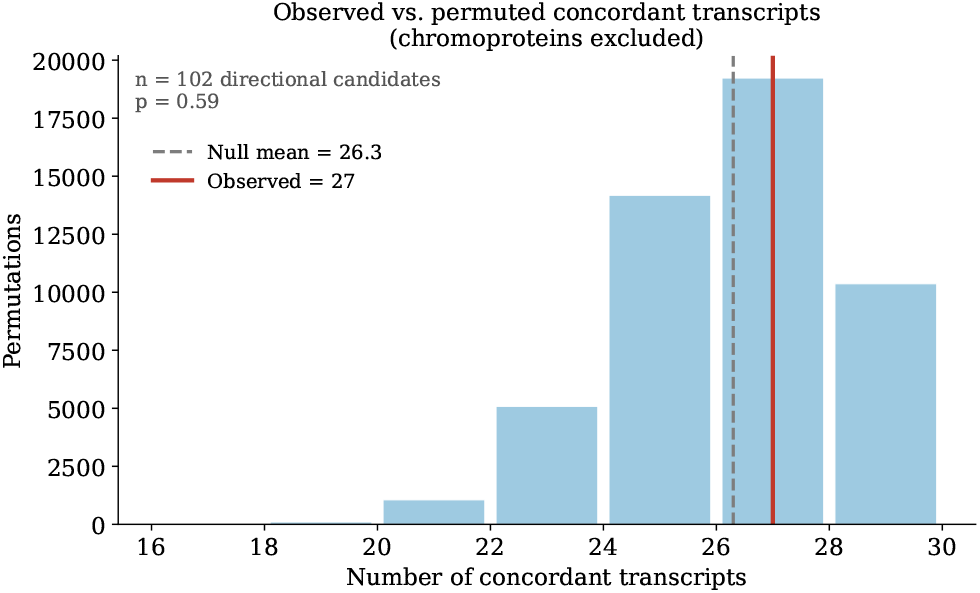
Expression direction does not match expected direction based on phylogenetic color class. A label-permutation significance test was perrformed comparing the observed number of direction-concordant candidates against the label-permutation null distribution. The observed value sits at the null mean, so pigment transcripts as a group do not change in the direction their color class predicts, and there is no convincing evidence that pigment transcript abundance is associated with phylogenetic color class.

The strictly monotonic subset gave the same result. Of the 56 strictly monotonic transcripts, 18 (32.1%) changed in the direction their color class predicted, and a label-permutation test restricted to this subset gave p = 0.78, again at chance.

We repeated the directional class prediction test on the most abundant transcripts, defined by peak host-normalized read count, on the rationale that transcript abundance may relate to the amount of pigment protein present and therefore to visible color. We tested the top 10 and top 50 transcripts by peak expression, as well as transcripts with more than 1,000 normalized counts. Restricting the analysis by abundance did not change the result: concordance remained between approximately 28% and 32% and did not exceed chance (Table 3).

Because the green-classified transcripts accounted for most of the disagreement between observed direction and class prediction, we examined all 96 in greater detail. Of these, 74 were higher in CO than in CG, splitting into 38 non-converting GFPs and 36 green-to-red photoconvertible EosFP-family proteins. Among the 18 of these 74 transcripts with peak expression of at least 1,000 normalized counts, 15 (83.3%) were photoconvertible. The six most abundant matched EosFP or d2EosFP references at 92 to 95% amino-acid identity, as shown in Table 4 and Figure 4. However, not all EosFPs examined in this study are more abundant in CO. Certain EosFPs, such as g39 as demonstrated by row 2 in Table 2, are more abundant in CG than CO, showing that while EosFPs may play a role in orange coloration, not all EosFP genes contribute to photoconversion-associated coloration and that EosFP-family transcripts do not behave uniformly.

**Table 1.** Leading warm pigment transcripts most highly expressed in the orange oral region (CO). Phylogenetic class gives the assigned color class and the nearest reference protein from FPbase. Values are host-normalized counts. CG is the green body, and CI is the intermediate transition region.

| Gene | Phylogenetic class | CG (green) | CI | CO (orange) |
|---|---|---|---|---|
| g65121 | OFP, melerfp-type | 2,035 | 5,868 | 37,859 |
| g26499 | chromoprotein, lea-type | 641 | 2,734 | 20,043 |
| g71578 | chromoprotein, lea-type | 532 | 1,581 | 13,205 |
| g94096 | chromoprotein, merufp-type | 635 | 1,972 | 12,788 |
| g45518 | chromoprotein, lea-type | 397 | 1,219 | 9,954 |
| g125134 | OFP, melerfp-type | 238 | 556 | 4,269 |
| g38342 | chromoprotein, mcfp506-type | 108 | 346 | 2,394 |
| g53223 | chromoprotein, echifp-type | 1,439 | 1,194 | 1,595 |
| g982 | chromoprotein, lea-type | 23 | 101 | 626 |
| g29516 | chromoprotein, mcfp506-type | 33 | 166 | 495 |

**Table 2.**
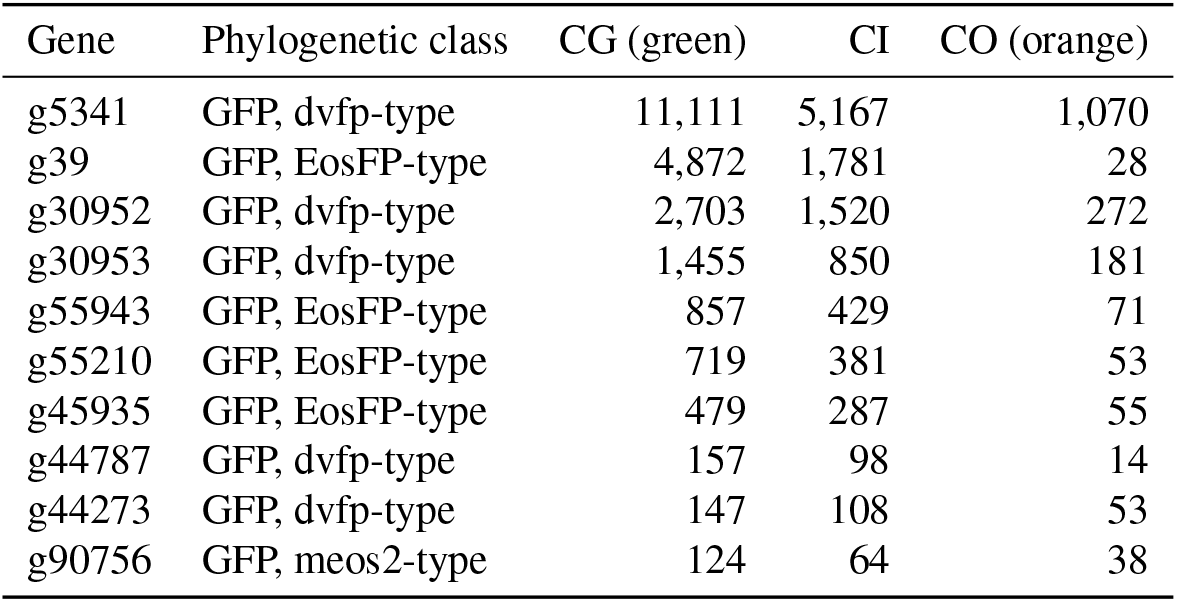
Leading green fluorescent protein (GFP) class transcripts most highly expressed in the green body (CG). Phylogenetic class gives the assigned color class followed by the nearest reference protein from FPbase that defines the subtype. Values are host-normalized counts. CI is the intermediate transition region, and CO is the orange oral region.

**Table 3.** Direction of expression change versus phylogenetic color-class prediction. A transcript is concordant if its change from the green body (CG) to the orange oral region (CO) matches its class prediction: GFP and CFP transcripts are predicted to decline toward CO, OFP and RFP to rise. The concordant column gives the count and percentage of n. P-values are from a label-permutation test shuffling class labels across transcripts with observed directions held fixed. Concordance does not exceed chance in any subset.

| Subset | *n* | concordant | *p* |
|---|---|---|---|
| Whole catalog (permutation) | 102 | 27 (26.5%) | 0.59 |
| Strictly monotonic (permutation) | 56 | 18 (32.1%) | 0.78 |
| Top-10 most abundant | 10 | 3 (30%) | 0.80 |
| Top-50 most abundant | 50 | 16 (32%) | 0.79 |
| Peak ≥ 1000 counts | 25 | 7 (28%) | 0.63 |
| Per class: GFP (predicted decrease) | 96 | 22 (23%) | — |
| Per class: OFP (predicted increase) | 4 | 3 (75%) | — |
| Per class: RFP (predicted increase) | 2 | 2 (100%) | — |

**Table 4.**
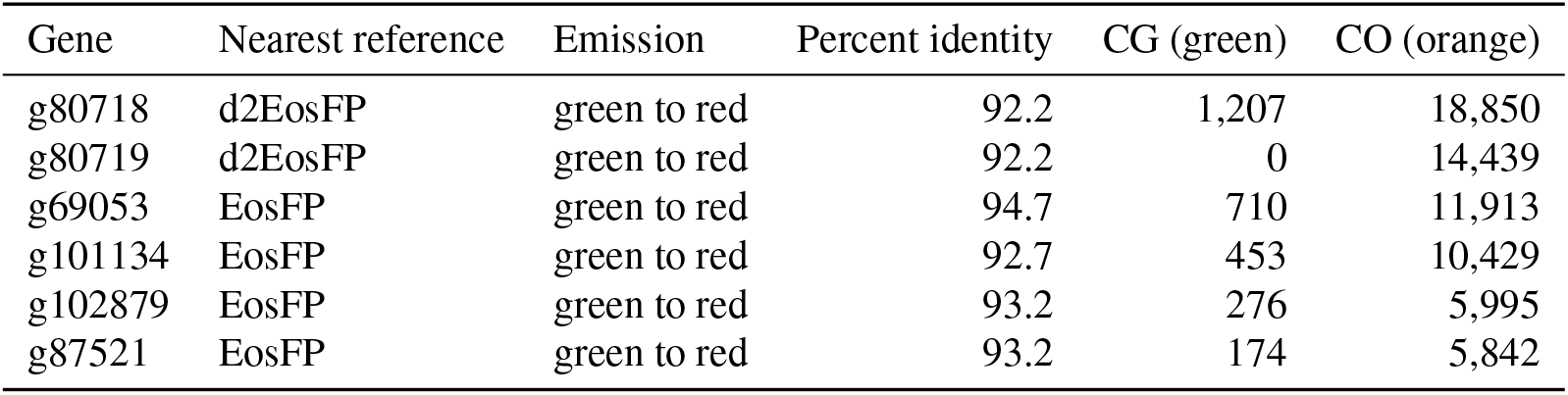
The six most abundant green-classified transcripts in CO are all EosFP- or d2EosFP-family photoconvertible proteins. Values are host-normalized counts. Identity is amino-acid percent to the nearest reference.

| Gene | Nearest reference | Emission | Percent identity | CG (green) | CO (orange) |
|---|---|---|---|---|---|
| g80718 | d2EosFP | green to red | 92.2 | 1,207 | 18,850 |
| g80719 | d2EosFP | green to red | 92.2 | 0 | 14,439 |
| g69053 | EosFP | green to red | 94.7 | 710 | 11,913 |
| g101134 | EosFP | green to red | 92.7 | 453 | 10,429 |
| g102879 | EosFP | green to red | 93.2 | 276 | 5,995 |
| g87521 | EosFP | green to red | 93.2 | 174 | 5,842 |

**Figure 4.**
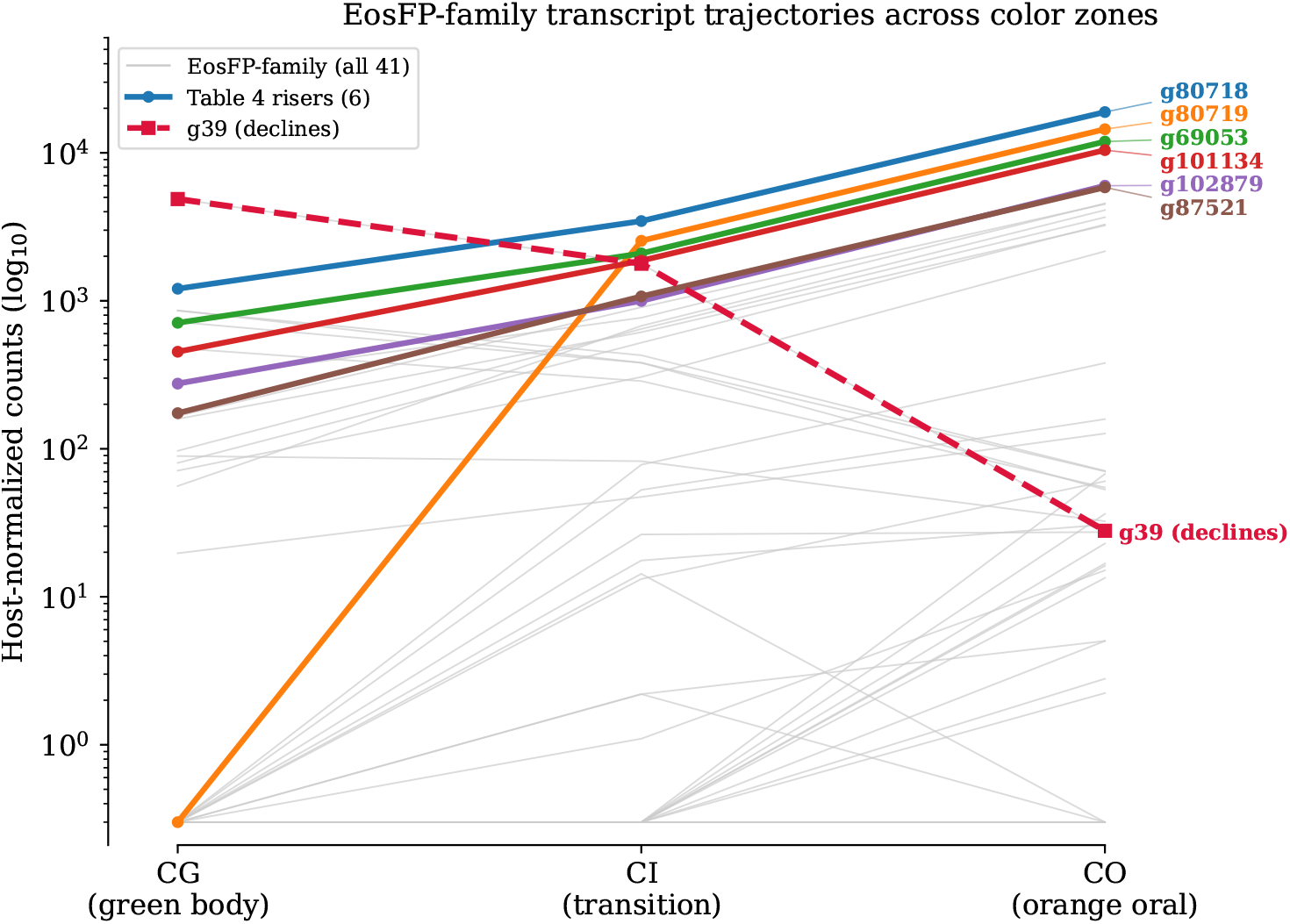
Most green-to-red photoconvertible EosFP-family transcripts are more abundant in the orange oral region, but the family is not uniform. All 41 transcripts whose nearest reference is an EosFP-family protein are plotted. Colored lines show the six most abundant green-classified transcripts that are highest in CO relative to CG (the same six transcripts as Table 4), each matched at 92 to 95% amino-acid identity to EosFP or d2EosFP references, along with g39, an EosFP-type transcript that is instead most abundant in the green body, shown for contrast. Thin gray lines are the remaining 34 EosFP-family transcripts. Values are host-normalized counts.

These findings indicate that phylogenetic color class is an unreliable predictor of a pigment gene’s contribution to visible color. Class describes the color a protein is determined by its sequence to emit, but for photoconvertible proteins, that describes only the unconverted state; the color in the tissue depends instead on the light the protein has received. Color in this system is therefore not fully specified by sequence. Because the green class is both the largest and the one containing these converters, aggregating expression by color class obscures the loci that could actually produce the observed coloration.

The symbiont read fraction also differed between the three color regions. As demonstrated in Figure 5, the symbiont read fraction declined from 68% in CG to 62% in CI and 32% in CO. Host read fraction rose correspondingly, from 20% to 25% to 48%. The two fractions do not sum to 100% in any region because the remaining mapped reads fall on unresolved transcripts, which were assigned to neither the host nor the symbiont set. CG has the highest symbiont read fraction and CO the lowest. This observation raises the possibility that algal abundance, rather than host pigment expression, makes up a portion of the reason behind the color difference. However, host-only normalization corrected for this difference, and the pigment-expression patterns remained, indicating that they are not an artifact of the differing symbiont read fractions between libraries, as shown in Figure 5B. Without correction, the distribution of green-to-orange change per transcript is shifted toward positive values across the host transcriptome, an artifact of the differing symbiont share between libraries. Host-only normalization re-centers the distribution at no change, removing approximately a 2.4-fold inflation.

**Figure 5.**
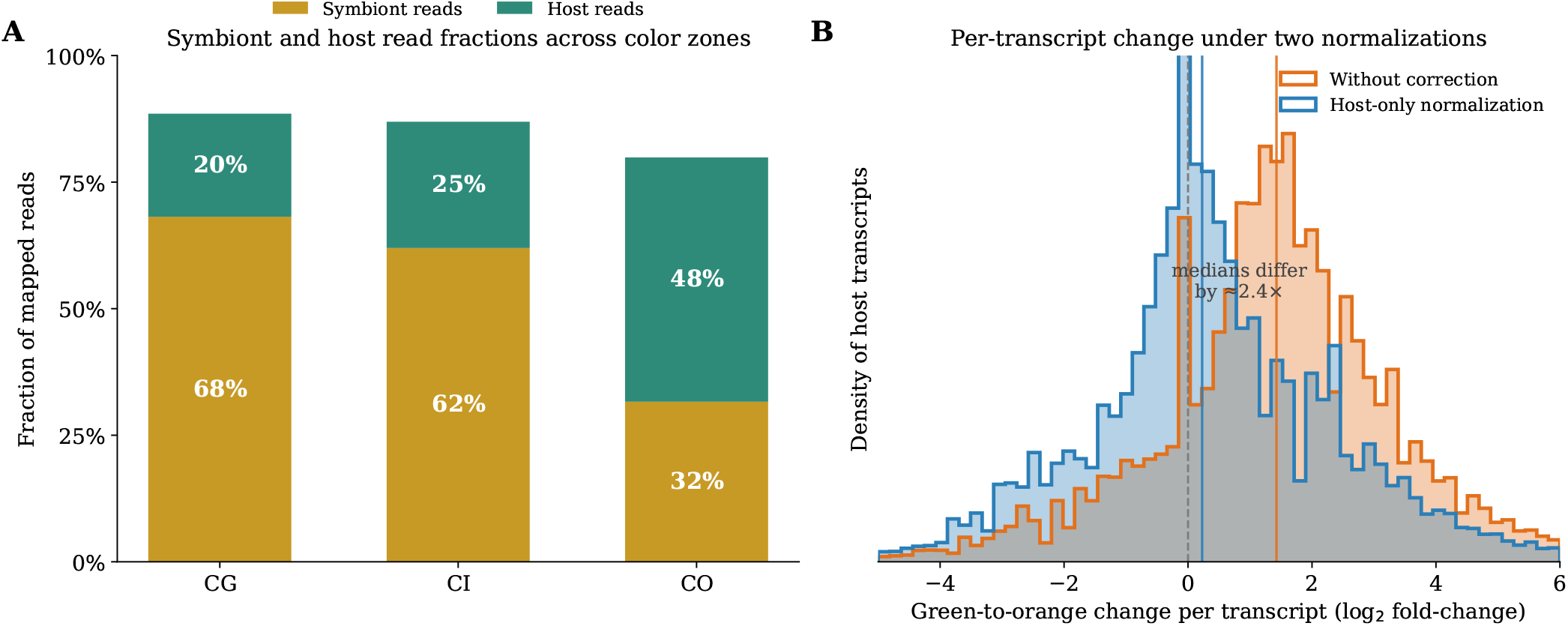
Correcting for algal read share changes what the data show. (A) In the green body and transition region, most reads come from the algal symbionts rather than the coral. The algal share drops sharply across the colony, from 68% in the green body to 32% in the orange region. Remaining reads fall on transcripts we could not assign to either partner. (B) This creates a problem. Standard normalization assumes libraries are comparable, so the heavy algal load in the green body makes coral transcripts there look artificially low. The result is that almost every coral gene appears higher in the orange region, whether or not it actually is. Computing the normalization from coral transcripts only fixes this. The orange curve is the uncorrected data, shifted right by roughly 2.4-fold. The blue curve is the corrected data, centered on no change.

## Discussion

Within one chalice coral colony, the green body and the orange oral region express many of the same pigment genes but at different levels. Because all regions of the colony share the same genetic background, their color differences are more likely to reflect differences in pigment-gene expression and possibly protein photoconversion. CO is associated with two coordinated transcriptional signals. The first is the up-regulation of distinct orange and chromoprotein transcripts relative to CG, which supports Hypothesis 2 and is the most direct contributor to the orange color. The second is the strong up-regulation of abundant green-to-red photoconvertible EosFP-family proteins in the same tissue, which supports Hypothesis 4.

Hypothesis 3 was partly supported. The symbiont share of the transcriptome does covary with color region, declining from 68% in CG to 32% in CO, but it does not account for the pigment signal, which persists after host-only normalization. The symbiont read fraction is therefore a real regional difference rather than a sufficient explanation for the color difference.

Expression across some of the pigment list does not follow phylogenetic color class because 77% of the green-classified transcripts found in CO are higher in CO than in CG. By nearest-reference identity, most of the abundant green-classified transcripts in CO are photoconvertible EosFP-family proteins. Many of these proteins were more highly expressed in the orange tissue, which suggests that photoconversion does contribute to the orange-red color. Similarly, Bollati et al. showed in mesophotic corals that partially photo-converted pcRFP tetramers support efficient intra-tetrameric FRET from the unconverted green chromophore to the photo-converted red chromophore, converting blue light into orange-red emission [12]. The closest matching reference proteins, EosFP and its variants, come from *Lobophyllia hemprichii* [9], a coral in the same family to which our specimen is tentatively assigned [26]. This supports the idea that these transcripts derive from coral loci.

## Limitations

The interpretation of this study has three limitations.

First, genetic data cannot reveal the physical spectral state of the photoconvertible proteins. High expression of a photoconvertible protein in orange tissue is consistent with photoconversion, but it does not imply that the protein is in its converted red state in the living tissue. A highly abundant EosFP, such as g39 in CG, may have been in its unconverted state of green or may have been in a realized state of red at the time of sampling the coral. Therefore, in principle, the orange color could arise wholly from the co-elevated chromoproteins and OFPs, while the photoconvertible protein remains green. Distinguishing these possibilities requires spectral or protein-level evidence, not transcript counts. Because the assembly is reference-free, individual transcripts cannot be confirmed as separate genomic loci; distinct contigs may represent isoforms or variants of a single gene. The qPCR validation proposed in the suggested further research section could help resolve this. Second, the two mechanisms, increased expression of chromoproteins and OFPs versus green-to-red protein photoconversion, are not mutually exclusive, and the study design cannot apportion their relative contributions. The orange region shows higher expression of both chromoprotein and OFP transcripts and green-to-red photoconvertible proteins.

Third, since only one colony was sampled, with a single library per region, the study lacks replication, constraining the scope of our conclusions. Within this colony, the expression differences among CG, CI, and CO are directly observed, rather than caused by differences in symbiont reads, because pigment signals persist after host-only normalization. However, because only one colony was studied, it is uncertain whether these patterns apply to other colonies or whether individual transcript changes are statistically significant at the population level.

### Suggested further research

Spectral confirmation of photoconversion in the EosFP-family proteins is needed. This step can potentially be completed through emission spectroscopy. The primary goal here is to test whether the orange tissue contains red-converted EosFP-family protein or whether the orange emission is instead produced by the co-elevated chromoproteins and OFPs. In parallel, this study should be biologically replicated, using five or more colonies with distinct color regions, to further verify the observed findings in this pilot study. The leading transcripts should also be validated by qPCR. These include the photoconvertible risers such as the g80718 d2EosFP-type transcript, the warm transcripts g65121 and g26499, and the body-enriched GFPs g5341 and g39.

## Conclusion

A single chalice coral colony shows different colors in its body and oral region because pigment genes are regulated differently across regions. The orange oral region is associated with two signals: 1) an up-regulation of distinct orange and chromoprotein transcripts; 2) a strong up-regulation of green- to-red photoconvertible EosFP-family proteins. Expression across the parts of the pigment list does not follow phylogenetic color class. For example, most green-classified transcripts were more abundant in the orange oral region than in the green body, in large part owing to the presence of photoconvertible proteins. The symbiont read fraction co-varies with color but does not solely explain it, because host-only normalization corrects for the differing symbiont read fractions between libraries and the pigment signals persist. As a reference-genome-free pilot study of a single colony, this work identifies a testable role for photoconversion in orange coloration. It also provides a reusable workflow that future studies can apply with biological replication and spectral confirmation added.

## Data availability

Raw RNA-seq reads are available in the NCBI Sequence Read Archive under BioProject PRJNA1559177. The de novo transcriptome assembly, the EosFP-like and pigment candidate sequences, the alignment, the phylogenetic tree, and the expression tables are available on Zenodo (https://doi.org/10.5281/zenodo.24094973).

